# A protein phosphatase network controls the temporal and spatial dynamics of differentiation commitment in human epidermis

**DOI:** 10.1101/125765

**Authors:** Ajay Mishra, Angela Oliveira Pisco, Benedicte Oules, Tony Ly, Kifayathullah Liakath-Ali, Gernot Walko, Priyalakshmi Viswanathan, Jagdeesh Nijjher, Sara-Jane Dunn, Angus I. Lamond, Fiona M. Watt

**Affiliations:** King’s College London, Centre for Stem Cells and Regenerative Medicine, 28th Floor, Tower Wing, Guy’s Hospital, Great Maze Pond, London SE1 9RT, UK.; Centre for Gene Regulation and Expression, School of Life Sciences, University of Dundee, Dow Street, Dundee, DD1 5EH, UK.; Microsoft Research, Cambridge CB1 2FB, UK; Wellcome Trust - Medical Research Council Cambridge Stem Cell Institute, University of Cambridge, Cambridge, CB2 1QR, UK

## Abstract

Epidermal homeostasis depends on a balance between stem cell renewal and terminal differentiation. The transition between the two cell states, termed commitment, is poorly understood. Here we characterise commitment by integrating transcriptomic and proteomic data from disaggregated primary human keratinocytes held in suspension for up to 12h to induce differentiation. We find that cell detachment induces a network of protein phosphatases. The pro-commitment phosphatases – including DUSP6, PPTC7, PTPN1, PTPN13 and PPP3CA – promote differentiation by negatively regulating ERK MAPK and positively regulating AP1 transcription factors. Their activity is antagonised by concomitant upregulation of DUSP10. Boolean network modelling of phosphatase interactions identifies commitment as an inherently unstable biological switch between the stem and differentiated cell states. Furthermore, phosphatase expression is spatially regulated both in vivo and in vitro. We conclude that an auto-regulatory phosphatase network maintains epidermal homeostasis by controlling the onset and duration of commitment.

## Introduction

Commitment is a transient state during which a cell becomes restricted to a particular differentiated fate. Under physiological conditions commitment is typically irreversible and involves selecting one differentiation pathway at the expense of others (Nimmo et al., 2015). While commitment is a well-defined concept in developmental biology, it is still poorly understood in the context of adult tissues (Simons and Clevers, 2011; Semrau and van Oudenaarden, 2015; Nimmo et al., 2015). This is because end-point analysis fails to capture dynamic changes in cell state, and rapid cell state transitions can depend on post-translational events, such as protein phosphorylation and dephosphorylation (Avraham and Yarden, 2011). For these reasons a systems approach to analyse commitment is required.

We set out to examine commitment in human interfollicular epidermis, which is a multilayered epithelium formed by keratinocytes comprising the outer covering of the skin (Watt, 2014). The stem cell compartment lies in the basal layer, attached to an underlying basement membrane, and cells that leave the basal layer undergo a process of terminal differentiation as they move through the suprabasal layers. In the final stage of terminal differentiation the cell nucleus and cytoplasmic organelles are lost and cells assemble an insoluble barrier, called the cornified envelope, which is formed of transglutaminase cross-linked proteins and lipids (Watt, 2014). Cells can commit to differentiate at any phase of the cell cycle, and upon commitment they are refractory to extracellular matrix (ECM)-mediated inhibition of differentiation (Adams and Watt, 1989).

Although there are currently no markers of commitment, we have previously used suspension-induced differentiation of disaggregated human keratinocytes in methylcellulose-containing medium (Adams and Watt, 1989) to define its timing, based on loss of clonal growth ability. From those studies, we have determined that commitment begins at approximately 4h and that terminal differentiation is initiated by 8h in suspension. We therefore used this simple experimental model to explore the nature of commitment in human keratinocytes.

## Results and Discussion

Since keratinocytes increase in size as they differentiate (Adams and Watt, 1989), we enriched for undifferentiated cells by a filtration step prior to placing them in suspension (Fig. 1a). By determining when cells recovered from suspension could no longer resume clonal growth on replating (Fig. 1b; Extended Data Fig. 1a), we confirmed that there is a marked drop in colony forming ability between 4 and 8 hours. This correlates with an increase in the proportion of cells expressing the terminal differentiation markers involucrin (IVL) and transglutaminase 1 (TGM1) (Fig. 1c, d; Extended Data Fig. 1b) and downregulation of genes that are expressed in the basal layer of the epidermis, including α6 integrin (ITGα6) and TP63 (Fig. 1d).

**Figure 1.**
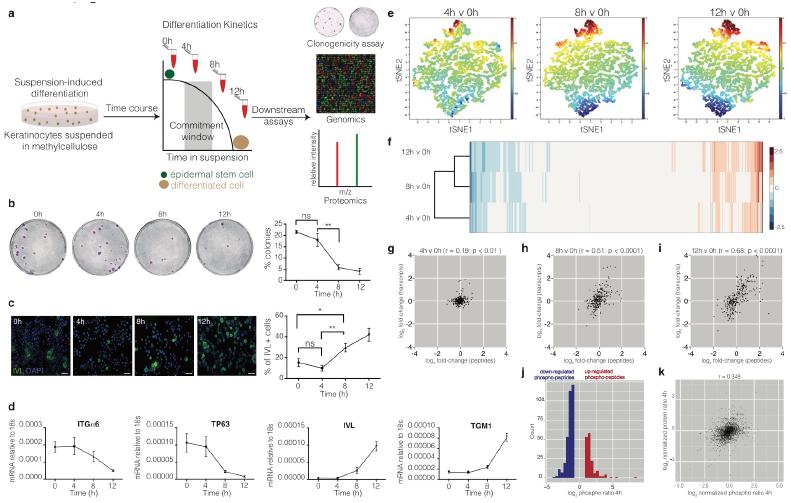
Genomic and proteomic analysis identifies protein dephosphorylation events that correlate with commitment. **a.** Schematic of experimental design. **b.** Colony formation by cells harvested from suspension at different times. Representative dishes are shown together with % colony formation (n=2 independent experiments, n= 3 dishes per condition per experiment; *p*-values calculated by Tukey’s multiple comparison test). **c.** Cells isolated from suspension at different time points were labelled with anti-involucrin (IVL) antibody (green) and DAPI as nuclear counterstain (blue). IVL-positive cells were counted using ImageJ (*n*=3 independent cultures; *p*-value calculated by 2-tailed t-test). Scale bars: 50μm **d.** RT-qPCR quantification of ITGα6, TP63, IVL and TGM1 mRNA levels (relative to 18s expression) (*n* = 3 independent cultures). **e.** t-SNE analysis of genome-wide transcript expression by keratinocytes placed in suspension for different times. **f.** Heatmap representing hierarchical clustering of differentially expressed proteins (*p* < 0.05). **g-i.** Dot plots correlating expression of significantly differentially expressed peptides (*p* < 0.05) that change two fold relative to 0h in at least one of the time points, with their corresponding differentially expressed transcripts. Pearson correlations (*r*) are indicated. **j.** Histogram of normalised SILAC ratios corresponding to high confidence phosphorylation sites that differ between 0 and 4h. **k.** Scatter plot correlating log_2_ normalized SILAC ratios for total protein changes (y-axis) with log_2_ phospho-peptide ratios (x-axis) between 0 and 4h. (**b, c)** \**p* < 0.05; \*\**p* < 0.01; ns = non-significant).

We next collected keratinocytes after 4, 8 and 12h in suspension and performed genome-wide transcript profiling using Illumina based microarrays and proteome-wide peptide analysis by SILAC-Mass-Spectrometry (MS) (Fig.1e, f; Extended Data Figs. 1c-f; Extended Data Tables 1, 2). Keratinocytes collected immediately after trypsinization served as the 0h control. When comparing the starting (0h) cell population with cells suspended for 4, 8 or 12h, t-SNE analysis of the gene expression data indicated that the 4h cell state was distinct from the 8 and 12h cell states (Fig. 1e). Unsupervised hierarchical clustering of differentially expressed genes (Extended Data Fig. 1c) or proteins (Fig. 1f) also indicated that the 4h sample clustered separately from the 8 and 12h samples. GO term enrichment analysis of differentially expressed transcripts or proteins showed enrichment of terms associated with epidermal differentiation at 8h and 12h (Extended Data Figs. 1e, f) (Sen et al., 2010; Mulder et al., 2012), which is consistent with the drop in clonogenicity seen at these time points.

When we compared the significantly differentially expressed proteins (*p*-value <0.05) that changed ≥2 fold at one or more time points with their corresponding transcripts (Fig. 1g-i), there was a moderately positive correlation at 8h and 12 h (Pearson correlations of 0.51 and 0.68, respectively), consistent with the correlation between bulk mRNA and protein levels seen in previous studies of mammalian cells (Schwanhäusser et al., 2011; Ly et al., 2014). However, at 4h transcripts and proteins were only weakly correlated (Pearson correlation 0.19, Fig. 1g).

The poor correlation between protein and transcript levels at 4h suggested a potential role for post-transcriptional mechanisms in regulating commitment. To investigate this we performed unbiased SILAC-MS based phospho-proteomic analysis. SILAC-labelled peptides isolated from cells at 0, 4 and 8h time points were enriched for phosphopeptides using HILIC prefractionation and titanium dioxide affinity chromatography (Extended Data Fig. 1d; Extended Data Tables 3, 4). Over 3,500 high confidence phosphorylation sites were identified with an Andromeda search score >= 30 and quantified at both the 4 and 8h timepoints. At 4h, approximately two thirds of the changes involved dephosphorylation (Fig. 1j). These dephosphorylation events could not be attributed to decreases in protein abundance, as shown by the discordance between total protein abundance and changes in protein phosphorylation (Fig. 1k).

Analysis of the integrated proteomic and genomic datasets revealed 45 phosphatases that were differentially expressed between 0 and 4h, 20 of which were upregulated (Fig. 2a, b). Interrogation of published datasets revealed that these phosphatases were also dynamically expressed during Calcium-induced stratification of human keratinocytes (Hopkin et al., 2012) and differentiation of reconstituted human epidermis (Lopez-Pajares et al., 2015) (Extended Data Fig. 2a, b).

**Figure 2.**
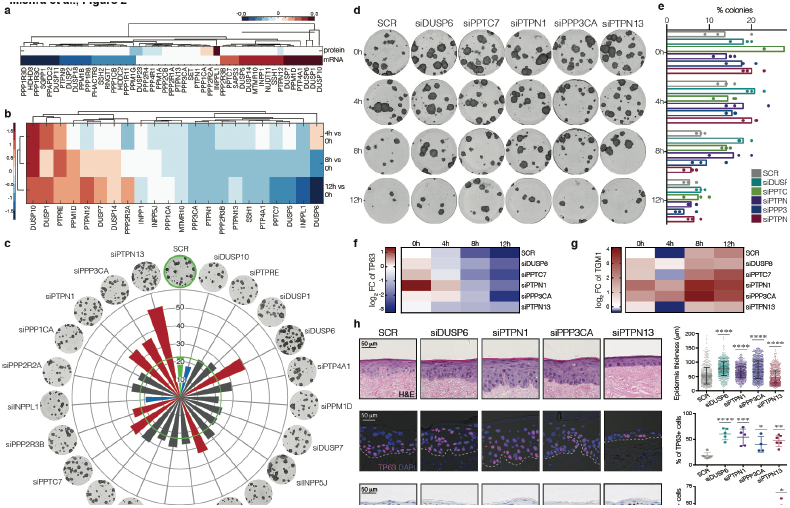
Functional screen identifies candidate phosphatases that regulate commitment. **a.** Heatmap representing the 47 phosphatases differentially expressed at transcript and/or protein level at 4h vs. 0h. **b.** Heatmap showing differential expression of those phosphatases that are upregulated at 4h in the microarray dataset at 4, 8 and 12h relative to 0h. **c.** Effect of knocking down the 21 phosphatases identified in (**b)** on clonal growth of keratinocytes. INPP5J was chosen as a control because it did not change in the microarray dataset. Values plotted are average *%* clonal growth in n=3 independent screens with n=3 independent cultures per screen. Green: SCR control. Red, blue: phosphatases with statistically significant effects on colony formation are shown (red: increase; blue: decrease). Grey: no statistically significant effect. **d, e.** Effect of knockdowns on clonal growth after 0h, 4h, 8h or 12h in suspension. (**d**) Representative dishes. **(e)** Mean % colony formation and individual values (*n*=*3* independent transfections). **f, g.** RT-qPCR quantification of TP63 (**f**) and TGM1 (**g**) mRNA levels (relative to 18s expression) in the same conditions as in **(d)**. *n*=3 independent transfections. **h.** Epidermal reconstitution assay following knockdown of DUSP6, PTPN1, PPP3CA or PTPN13. *n*=2 independent transfections. Top row shows representative H&E images. Epidermal thickness was quantified in multiple fields from 8 sections per replicate ± SD relative to scrambled control (SCR). Middle row shows staining for TP63 (pink) with DAPI nuclear counterstain (blue). % DAPI labelled nuclei that were TP63+ was quantified in *n*=2-3 fields per replicate. Bottom row shows staining for Ki67 (brown) with haematoxylin counterstain (blue). % Ki67+ nuclei was quantified in *n*=3-6 fields per replicate. Error bars represent mean ± s.d. *p*-values were calculated using one-way ANOVA with Dunnett’s multiple comparisons test (*p < 0.05; *\*\*p* < 0.01; *\*\*\*\*p* < 0.0001; ns = non-significant).

To examine the effects on keratinocyte self-renewal of knocking down each of the 20 phosphatases that were upregulated on commitment in suspension, we transfected primary human keratinocytes with SMART pool siRNAs and measured colony formation in culture (Fig. 2c; Extended Data Fig. 3a, b). As a control we also included INPP5J, which did not change in expression during suspension-induced differentiation. Live cell imaging was used to monitor cell growth for three days post-transfection (Extended Data Fig. 3b). Knocking down seven of the phosphatases significantly increased clonal growth (Fig 2c; Extended Data Fig. 3d), with five phosphatases – DUSP6, PPTC7, PTPN1, PTPN13 and PPP3CA - having the most pronounced effect (p-value <0.001). Silencing of these phosphatases also increased colony size and delayed the loss of colony forming ability in suspension (Fig. 2d, e; Extended Data Fig. 3d). Consistent with these findings, phosphatase knockdown delayed the decline in TP63 and increase in TGM1 levels during suspension induced differentiation (Fig. 2f, g; Extended Data Table 5). Cumulatively, their effects on keratinocyte self-renewal and differentiation suggest that these are pro-commitment protein phosphatases.

The effects of knocking down DUSP10 differed from those of knocking down DUSP6, PPTC7, PTPN1, PTPN13 and PPP3CA. DUSP10 knockdown significantly reduced clonogenicity (p-value<0.001; Fig. 2c; Extended Data Fig. 3a) and, in contrast to the other phosphatases, decreased the growth rate of keratinocytes (Extended Data Fig. 3b, e). In addition, whereas expression of DUSP6, PPTC7, PTPN1, PTPN13 and PPP3CA declined by 8h in suspension, DUSP10 expression remained high (Fig. 2b). Thus DUSP10 may serve to antagonise commitment.

To examine the effects of knocking down the pro-commitment phosphatases on the ability of keratinocytes to reconstitute a multi-layered epithelium, we seeded cells on de-epidermised human dermis and cultured them at the air-medium interface for three weeks (Fig. 2h; Extended Data Fig. 3f). Knockdown of DUSP6, PTPN1, PPP3CA and PTPN13 did not prevent cells from undergoing terminal differentiation, as evidenced by suprabasal expression of involucrin and accumulation of cornified cells. However, the proportion of TP63-positive cells was increased. Knockdown of DUSP6, PTPN1 or PPP3CA increased epidermal thickness without increasing the proportion of Ki67-positive, proliferative cells. Conversely, PTPN13 knockdown led to an increase in Ki67-positive cells and a reduction in epidermal thickness, which could reflect an increased rate of transit through the epidermal layers. In addition, Ki67 and TP63-positive cells were no longer confined to the basal cell layer of epidermis reconstituted following phosphatase knockdown, but were also present throughout the viable suprabasal layers. Thus suprabasal cells simultaneously expressed basal (TP63, Ki67) and suprabasal (IVL) markers, indicating that the transition from stem cells to differentiated cells had been disturbed.

To identify the signalling networks affected by upregulation of phosphatases during commitment we performed GO analysis of ranked peptides that were dephosphorylated at 4h. The top enriched pathways were ErbB1 signalling, adherens junctions, insulin signalling and MAPK signalling (Fig. 3a). Several of the proteins we identified are components of more than one pathway (Fig. 3b) and all have been reported previously to regulate epidermal differentiation (Connelly et al., 2010; Haase et al., 2001; Kolev et al., 2008; Scholl et al., 2012; Trappmann et al., 2012). Furthermore, constitutive activation of ERK delays suspension-induced differentiation (Haase et al., 2001).

**Figure 3.**
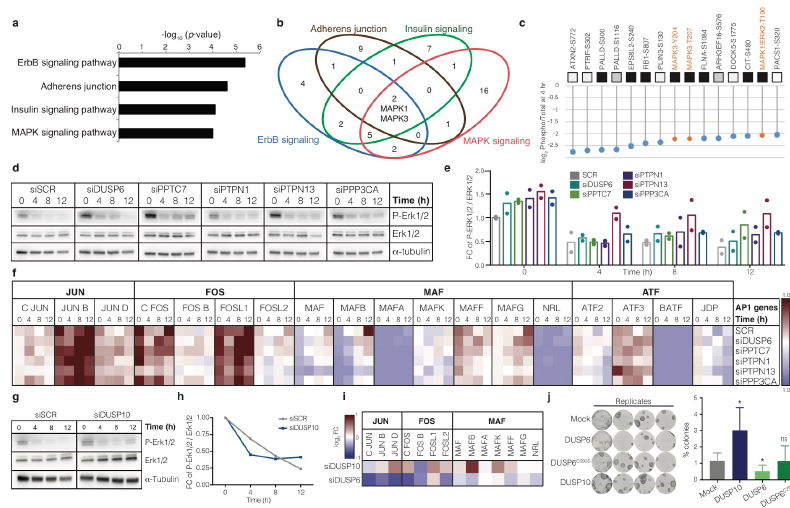
Pro-commitment phosphatases regulate MAPK signalling and AP1 transcription factors. **a.** Gene Ontology (GO) term enrichment analysis of ranked peptides dephosphorylated at 4h. **b.** Venn diagram showing intersection of signalling pathways regulated at 4h. **c.** Top 15 dephosphorylated peptide sites at 4h, showing ratio between change in phospho-peptides and change in total protein. Highlighted in orange are the MAPK sites. **d, e.** Western blots **(e)** showing phospho-ERK1/2 and total ERK1/2 in cells harvested after 0, 4, 8 and 12h in suspension. α-tubulin: loading control. Quantification of phospho-ERK relative to total ERK in n = 2 blots is shown in (**e**). Mean and individual values are plotted. **f.** Heatmap represents mRNA expression (relative to 18s RNA) of AP1 transcription factor superfamily members during suspension-induced differentiation post-phosphatase knockdown (*n*=*3;* values plotted are means of 3 independent transfections). See Extended Data Table 6 for *p*-values generated by 2-way ANOVA. **g-h.** Western blot of phospho-ERK1/2 and total ERK1/2 in suspended cells following DUSP10 knockdown. siSCR, loading controls and quantitation are shown. **i.** Heatmap represents mRNA expression (relative to 18s mRNA) of JUN, FOS and MAF family members after DUSP6 and DUSP10 knockdown (values plotted are means of 3 independent transfections normalised against scrambled control; see Extended Data Table 7 for *p*-values generated by 2-way ANOVA). **j.** Clonal growth (representative dishes and quantification) following doxycycline-induced over-expression of DUSP10, DUSP6 and mutant DUSP6^C293S^ in primary keratinocytes (*n*=3 independent cultures). *p*-values were calculated using one-way ANOVA with Dunn’s multiple comparisons test (\**p* < 0.05; ns = non-significant).

We next ranked protein phosphorylation sites according to the log_2_-fold decrease at 4 hours, plotting the ratio between the change in phosphorylation site and the change in total protein (Extended Data Table 5). To specifically identify dephosphorylation events, we only considered proteins that increased in abundance by more than 0.5 in log_2_ space while phosphorylation levels that remained constant were excluded from the ranking. Consistent with the predicted dynamic interactions between signalling pathways (Fig. 3b), phosphorylation sites on MAPK1 (ERK2) and MAPK3 (ERK1) were identified in the top 15 most decreased sites (Fig. 3c). Other proteins in the top 15 included components or regulators of the cytoskeleton (FLNA), Rho signalling (DOCK5, ARHGEF16, CIT) and EGFR signalling (EPS8), again consistent with the GO terminology analysis.

We performed Western blotting to confirm that ERK1/2 activity was indeed modulated by suspension-induced terminal differentiation and by the candidate pro-commitment phosphatases (Fig. 3d, e). As reported previously, the level of phosphorylated ERK1/2 diminished with time in suspension (Janes et al., 2009) (Fig. 3d). Knockdown of the pro-commitment phosphatases (Extended Data Fig. 3c) resulted in higher levels of phosphorylated ERK1/2 relative to the scrambled control, both at 0h and at later time points (Fig. 3d, e). These effects are consistent with the known requirement for ERK MAPK activity to maintain keratinocytes in the stem cell compartment (Trappmann et al., 2012).

Transcriptional regulation of epidermal differentiation is mediated by the Activator Protein 1 (AP1) family of transcription factors (Eckert et al., 2013), which is the main effector of the MAPK and ErbB signalling cascades (Karin and Chang, 2011). Quantification of the levels of AP1 transcripts during suspension-induced terminal differentiation revealed that different AP1 factors changed with different kinetics, as reported previously (Gandarillas and Watt, 1995) (Fig. 3f). Notably, the level of several members of the MAF subfamily of AP1 factors (MAF, MAFB and MAFG) significantly increased during differentiation, consistent with recent evidence that they mediate the terminal differentiation programme in human keratinocytes (Lopez-Pajares et al., 2015). In line with these observations, knockdown of individual pro-commitment phosphatases reduced the induction of MAF AP1 factors in suspension (Fig. 3f; Extended Data Table 6). These experiments are consistent with a model whereby induction of phosphatases in committed keratinocytes causes dephosphorylation of ERK MAPK and prevents the increase in expression of AP1 transcription factors that execute the terminal differentiation programme.

To explore why the commitment phase of suspension-induced terminal differentiation was transient, we focused on DUSP10, which – like the pro-commitment phosphatases – was upregulated at 4h, yet had the opposite effect on clonal growth (Fig. 2c; Extended Data Fig. 3a, b, e). Unlike knockdown of DUSP6, PPTC7, PTPN1, PTPN13 or PPP3CA, knockdown of DUSP10 (Extended Data Fig. 4a, b) did not increase ERK1/2 activity (Fig. 3g). Although, consistent with its known selectivity for p38 MAPK (Caunt and Keyse, 2013), DUSP10 knockdown increased phospho-p38 at 0h, there was no effect at later times in suspension (Extended Data Fig. 4c, d). Again, in contrast to the pro-commitment phosphatases, DUSP10 knockdown increased expression of several AP1 transcription factors, including members of the JUN, FOS and MAF subfamilies (Fig. 3i; Extended Data Table 7).

We confirmed the antagonistic effects of DUSP6 and DUSP10 by overexpressing each phosphatase in human keratinocytes with Cumate or Doxycycline inducible vectors (Fig. 3j; Extended Data Fig. 4e-g). As predicted, whereas overexpression of DUSP10 increased colony formation, overexpression of DUSP6 reduced clonal growth (Fig. 3j). A dominant negative mutant of DUSP6 (C293S) lacking phosphatase activity (Okudela et al., 2009) had no effect (Fig. 3j; Extended Data Fig. 4e-g).

To examine whether the different phosphatases interacted genetically, we performed individual knockdowns of the 5 pro-commitment phosphatases and DUSP10 after 0, 4, 8 or 12h in suspension, and examined the effect on mRNA levels of the other phosphatases (Extended Data Fig. 5a; Extended Data Table 8). This led us to infer the regulatory networks depicted in Fig. 4a, where the node colours show fold-change relative to the 0h time-point. Arrows indicate positive effects on expression and T-bars show inhibitory effects. The analysis indicates a key role for DUSP6 at commitment time (4h), when DUSP6 expression is positively regulated by a self-amplifying loop as well as by all other phosphatases in the network. At all other time points the interactions between individual phosphatases are predominantly negative. Several phosphatases, including PTPN1 and PTPN13, show positive autoregulation at one or more time points. It is also notable that DUSP10 is negatively regulated by other phosphatases, except at 4h, when it is positively regulated by PTPN1 and by an autoregulatory loop (Fig. 4a).

**Figure 4.**
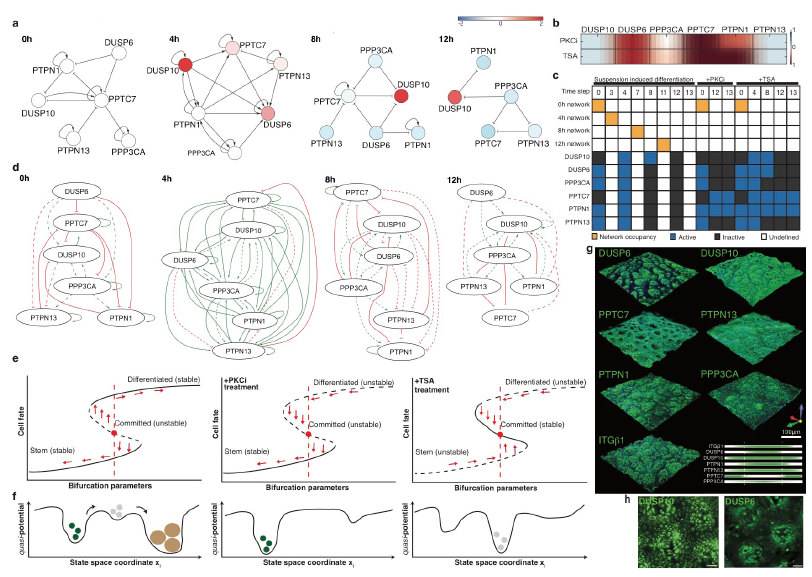
An autoregulatory network of phosphatases controls commitment. **a.** Colours represent log2 fold-change in phosphatase expression relative to 0h (values plotted are means of 3 independent experiments normalised against SCR control). **b.** Heatmap represents mRNA expression (relative to 18s mRNA) of individual phosphatases in cells treated in suspension for 12h with PKCi or TSA (values plotted are the means of 3 independent experiments normalised against vehicle-treated control). **c.** Boolean network iteration scheme. We defined 8 experimental constraints, 4 for suspension-induced differentiation in the absence of pharmacological inhibitors, 3 for TSA and 1 for PKCi treatment. For cells in the absence of drugs we imposed a switching scheme, whereby the system must change the representative network in order to achieve the expression constraints. **d.** Networks able to satisfy the model constraints of the Boolean formalism in (**c**) are depicted. Solid lines show interactions already calculated in (**a**), while dashed lines were inferred from Extended Data Fig. 5d, e. See also Extended Data Tables 9, 10. **e, f.** Representation of commitment as possible two saddle-node bifurcations in a direction xi of the state space for control cells or cells treated with PKCi or TSA. In the control both stem and differentiated cell states are stable (attractors), while commitment is an unstable state. Since the system is able to reach the expression constraint for PKCi at 12h, we hypothesize that on PKCi treatment, the only stable state is the stem state. On TSA treatment there is a mandatory switch from the 0h network but the 12h network cannot be reached at any time point; we therefore hypothesize that commitment becomes a stable state while the stem and differentiated cell states are unstable. **g.** 3D-volume rendered confocal images of wholemounts of human epidermis labelled with antibodies against commitment phosphatases or ITGβ1 (green) and counterstained with DAPI (blue). The distribution of each phosphatase relative to ITGβ1 is also shown graphically. **h.** 3D-volume rendered confocal images of primary keratinocytes cultured on PDMS topographic substrates and labelled with antibodies to DUSP10 and DUSP6. Scale bars: 100μm.

The changes in the phosphatase interaction network topology with time in suspension suggest that a biological switch occurs at 4h. The negative feedback loops predominating at 0h, 8h and 12h are known to result in stable phenotypes because the network is able to counteract additional inputs (Zeigler et al., 2000). However, at 4h all but one of the interactions are positive. Positive feedback loops lead to instability, because the network amplifies any inputs it receives. The concept of commitment as an unstable state is supported by the experimental evidence that it is transient.

To test the robustness of the network we examined the effects of treating keratinocytes with the histone deacetylase inhibitor Trichostatin A (TSA) and a Protein Kinase C inhibitor (PKCi). Both drugs blocked suspension-induced terminal differentiation, as measured by expression of IVL and TGM1, and prevented downregulation of ITGα6 (Extended Data Fig. 5a, b). However, cells treated with TSA still underwent commitment, as evaluated by loss of colony forming ability and downregulation of TP63, whereas those treated with the PKC inhibitor did not. At 12h in suspension both drugs reduced expression of DUSP10 and PTPN13 and increased expression of DUSP6 and PPTC7 relative to untreated cells; however, they differentially affected PP3CA and PTPN1 (Fig. 4b).

We tested whether the inferred network switch associated with commitment is mandatory, by employing a Boolean network abstraction to explore the dynamics of the network (Fig. 4c). Accordingly, we abstracted gene expression levels as ‘on’ or ‘off’ if their mean expression was respectively higher or lower than the average of all genes. We sought to confirm that the network topologies at each timepoint (Fig. 4a) were consistent with experimental observations of gene expression, by employing the automated reasoning approach encapsulated in the tool RE:SIN, as described by Shavit et al. (2016). This approach allowed us to determine whether the inferred networks could recapitulate the dynamic changes in gene expression. To this end, we defined a set of 3 experimental constraints, comprising suspension-induced differentiation at different time points and PKCi and TSA treatment (Fig. 4c).

We first found that the networks inferred by the genetic knockdown experiments could not satisfy these 3 experimental observations, suggesting that additional interactions exist between the phosphatases that were not revealed by these data. Therefore, we incorporated additional ‘possible’ interactions, to construct Abstract Boolean Networks (ABNs) for each stage. This formalism enables the investigation of putative interactions, as an ABN implicitly defines a set of concrete Boolean networks each with a unique topology (Yordanov et al., 2016). We included these possible interactions based on the effects of overexpressing DUSP6 and DUSP10, and by examining phosphatase protein levels (Extended Data Fig. 5; Extended Data Table 9).

By extending the ABN at each time point to incorporate possible phosphatase interactions, we found that the model constraints could be satisfied (Fig. 4d; Extended Data Table <u>10</u>). Crucially, we also tested whether these constraints could be satisfied by a single ABN, to examine whether the gene expression changes could be explained without employing network switching. To this end, we constructed an ABN in which all phosphatases were allowed to interact either positively or negatively. Using the automated reasoning approach encapsulated in the sister tool RE:IN, we deduced that a single Boolean network was unable to recapitulate the measured expression dynamics, and hence that the network must change with time.

We next tested whether network switching still occurs under PKCi and TSA treatment using the sets of concrete Boolean networks that we derived for each stage. In both cases, we found that we could not transit through the networks and respect the expected expression states, corroborating the experimental finding that terminal differentiation is blocked in these conditions. The PKCi phosphatase expression pattern at 12h was compatible with the phosphatase interaction network derived at 0h, supporting the conclusion that PKCi arrests cells in the stem cell compartment. In TSA treated cells the network must switch from that derived at 0h to that derived at 4h, and subsequently to that derived at 8h, or straight from 0h to 8h. Importantly, neither PKCi nor TSA treatment resulted in an expression pattern compatible with the network derived at 12h for untreated cells, consistent with the inhibition of differentiation.

Using dynamical systems terminology, we can describe epidermal differentiation as two saddle-node bifurcations (Fig. 4e, f). We start with a single minimum (μ_stem_), then pass through a first saddle-node bifurcation to have two minima (μ_committed*_), then the global minimum changes (μ_commited**_) and finally a second saddle-node bifurcation leads again to a single steady state (μ_differentiated_) (Zhang et al., 2012). The stem cell state and the terminally differentiated state emerge as stable states, while commitment is inherently unstable and serves as a biological switch.

We next examined whether the epidermal phosphatase network we identified was also subject to spatial regulation, since spatiotemporal coordination of stem cell behaviour contributes to epidermal homeostasis (Doupé and Jones, 2012). By wholemount labelling the basal layer of sheets of human epidermis (Jensen et al., 1999) we found that DUSP6, PTPN1 and PPP3CA were most highly expressed in cells with the highest levels of β1 integrins, which correspond to stem cell clusters (Jensen et al., 1999) (Fig. 4g). In contrast, PPTC7 was enriched in the integrin-low regions, while PTPN13 and DUSP10 were more uniformly expressed throughout the basal layer (Fig. 4g). The inverse relationship between the patterns of PPP3CA and PPTC7 expression is in good agreement with the network analysis demonstrating negative regulation of PPP3CA by PPTC7 in undifferentiated keratinocytes (0h; Fig. 4a).

The patterned distribution of phosphatases could be recreated in vitro by culturing keratinocytes on collagen-coated PDMS elastomer substrates that mimic the topographical features of the human epidermal-dermal interface (Viswanathan et al., 2016). We observed clusters of cells with high levels of DUPS6 on the tops of the features, where stem cells expressing high levels of β1 integrins accumulate (Viswanathan et al., 2016) (Fig. 4h). In contrast, DUSP10 was uniformly expressed regardless of cell position (Fig. 4h). These results indicate that the phosphatases are subject to spatial regulation that is independent of signals from cells in the underlying dermis.

In conclusion, we have shown that epidermal commitment is a biological switch controlled by a network of protein phosphatases that are regulated spatially and temporally. The key role of DUPS6 at commitment fits well with its upregulation by Serum Response Factor, which is known to control keratinocyte differentiation (Connelly et al., 2010) and the importance of DUSP6 in controlling the activation kinetics and dose-response behaviour of ERK MAPK signalling (Blüthgen et al., 2009). The involvement of multiple phosphatases may protect cells from undergoing premature terminal differentiation and is consistent with the ability of different external stimuli to trigger differentiation via different intracellular pathways (Watt, 2016). Furthermore, the upregulation of basal layer markers in the suprabasal epidermal layers on knockdown of pro-commitment phosphatases mimics features of psoriatic lesions in which ERK is known to be upregulated (Haase et al., 2001), leading us to speculate that commitment is deregulated in hyperproliferative skin conditions.

## Methods

### Cell culture

Primary human keratinocytes (strain km) were isolated from neonatal foreskin and cultured on mitotically inactivated 3T3-J2 cells in complete FAD medium, containing 1 part Ham’s F12, 3 parts Dulbecco’s modified eagle medium (DMEM), 10^−4^ M adenine, 10% (v/v) FBS, 0.5 μg ml^-1^ hydrocortisone, 5 μg ml^-1^ insulin, 10^−10^ M cholera toxin and 10 ng ml^-1^ EGF, as described previously (Gandarillas and Watt, 1995). Prior to suspension-induced differentiation in methylcellulose (Adams and Watt, 1989) we enriched for stem cells by filtering the disaggregated keratinocytes twice through a 40μm sterile membrane. For knockdown or overexpression experiments, keratinocytes were grown in Keratinocyte-SFM medium (Gibco) supplemented with 0.15 ng/ml EGF and 30 μm/ml BPE. All isoforms of PKC were inhibited using 5 μΜ GF 109203X (Tocris; at lower concentrations GF190203X preferentially inhibits the -α and -β isoforms) and histone deacetylase was inhibited with TSA (Sigma Aldrich). PDMS substrates that mimic the topography of the epidermal-dermal junction were generated as described previously (Viswanathan et al., 2016).

### Genome-wide expression profiling

Genome-wide expression profiling was performed using the Illumina BeadArray platform and standard protocols. Data were processed using R (ISBN 3-900051-07-0/ http://www.r-project.org/) or Genespring GX13.1 software. The gene expression data are deposited in the GEO databank (GSE73147).

### Computational analysis of gene expression datasets

Microarray initial processing and normalization were performed with BeadStudio software. BeadChip internal *p*-values (technical bead replicates) were used for filtering out genes significantly expressed above the background noise. To filter out genes with signals that were not significant, a *p*-value of 0.05 was used as the cut-off value and only genes with a *p*-value <0.05 in at least one sample passed the filter. From the original set of 34,685 gene targets, 23,356 targets met this criterion. The data were imported into GeneSpring v13, normalized using quantile normalization and the biological replicates averaged for subsequent analysis. We performed pairwise comparison between 0h and 4, 8 and 12h. Genes showing a fold change higher than 2 comparing to control (and both *p*-values were significant) were subjected to GO analysis with GeneSpring v13 (152 genes between 4h and 0h, 553 between 8 and 0h, 1136 genes between 12 and 0h). Hierarchical clustering based on Pearson’s uncentred distance was performed on time course gene expression data and the results presented as a heatmap.

### Network analysis

A 2-way ANOVA with multiple comparisons corrected using the Holm-Sidak test was used to identify the statistically significant effects of single phosphatase knockdown on the expression of the other phosphatases. The weight of each edge was calculated as the inverse p-value for the respective interaction. For simplification we kept only the significant links. The Boolean network analysis was performed using the RE:IN and RE:SIN software, which is designed to encode possible genetic interactions and behavioural constraints (Dunn et al., 2014; Shavit et al., 2016; Yordanov et al., 2016) A set of possible and definite interactions define an Abstract Boolean Network, which implicitly defines a set of concrete Boolean networks, with unique topologies defined by the presence or absence of the possible interactions. The software allows the user to seek the set of concrete Boolean network models that are consistent with a set of experimental constraints, and to use this constrained set to make predictions of untested behaviour. Experimental constraints are expected states along network trajectories, which are constructed from discretised gene expression patterns. The software is freely available to use, and can be accessed together with tutorials and FAQ at research.microsoft.com/rein.

### Generation and analysis of SILAC LC-MS/MS datasets

A pre-confluent keratinocyte culture was split into two separate cultures with FAD^-lysine-arginine^ medium (Sigma) differentially supplemented either with K^0^R^0^ or with K^8^R (stable isotopes of amino acids Lysine and Arginine; Cambridge Isotope Laboratories) (Ly et al., 2014). FCS used for SILAC medium was also depleted of Lysine and Arginine (Sigma). Cells were grown in SILAC medium for 5-6 days to reach 70 – 80% confluence and later harvested for downstream assays. Light labelled cells served as the 0h sample whereas heavy labelled cells were suspended in methylcellulose and harvested at 4, 8 and 12h. Cell extracts prepared from 0h cells were mixed individually with 4, 8 and 12h samples at a 1:1 ratio of total protein. The mixed samples were then subjected to mass spectrometry (MS).

For measurement of protein abundance changes by SILAC, lysates containing ∼80 μg protein were prepared in LDS sample buffer containing reducing agent (TCEP). Proteins were separated by electrophoresis on a 4-12% Bis-Tris NuPAGE gel, which was Coomassie-stained and cut into eight equally sized gel slices. Gel-embedded proteins were reduced with TCEP, alkylated with iodoacetamide, and trypsin-digested to release peptides. Peptides were extracted from gels using 50% acetonitrile (ACN) containing 5% formic acid (FA), dried and resuspended in 5% FA. Peptides were then analysed by LC-MS/MS on a Dionex RSLCnano coupled to a Q-Exactive Orbitrap classic instrument. Specifically, peptides were loaded onto a PepMap100 75μm × 2cm trap column, which was then brought in-line with a 75μm × 50cm PepMap-C18 column and eluted using a linear gradient over 220 min at a constant flow rate of 200 nl/min. The gradient composition was 5% to 35% B, where solvent A = 2% ACN + 0.1% FA and solvent B = 80% ACN + 0.1% FA. Peptides were eluted with a linear elution gradient (5% B to 35% B) over 220 min with a constant flow rate of 200 nl/min. An initial MS scan of 70,000 resolution was acquired, followed by data-dependent MS/MS by HCD on the top 10 most intense ions of ≥2+ charge state at 17,500 resolution.

For phosphoenrichment analysis, lysates containing ∼2 mg of protein were chloroform:methanol precipitated. Pellets were resuspended in 8M urea in digest buffer (0.1 M Tris pH 8 + 1 mM CaCl^2^), diluted to 4 M urea with additional digest buffer, and digested with Lys-C for 4h at 37°C. The digests were diluted to 1M urea and trypsin-digested overnight at 37°C. Digests were then acidified and desalted using SepPak-C18 vacuum cartridges. Desalted peptides were resuspended in mobile phase A for HILIC (80% ACN + 0.1% TFA). HILIC chromatography was performed using a Dionex Ultimate 3000 with a TSK Biosciences Amide-80 column (250 × 4.6 mm) (Di Palma et al., 2013; McNulty and Annan, 2008; Navarro et al., 2011). Peptides were eluted using an exponential gradient (80% B to 60% B) composed of A (above) and B (0.1% TFA) at 0.4 ml/min over 60 min. 16 fractions were collected from 25 min to 60 min. These fractions are enriched for hydrophilic peptides, including phosphopeptides. The fractions were dried before further phosphoenrichment by titanium dioxide (TiO_2_).

For TiO_2_ enrichment, HILIC fractions were resuspended in loading buffer (70% ACN + 0.3% lactic acid + 3% TFA). 1.25 mg of TiO_2_ (GL Sciences) was added to each fraction and incubated for 10 min to bind phosphopeptides. Beads were washed with loading buffer, and two wash buffers, composed of (1) 70% ACN + 3% TFA and (2) 20% ACN + 0.5% TFA. Phosphopeptides were eluted in two steps, first with 4% of ammonium hydroxide solution (28% w/w NH_3_) in water for 1h and again with 2.6% of ammonium hydroxide solution in 50% ACN overnight. Elutions were collected, dried, resuspended in 5% FA and analysed by LC-MS/MS. The LC-MS analysis was performed similarly to above with the following modifications: peptides were chromatographed on an EasySpray PepMap 75 μm × 50 cm column, and a ‘Top30’ method was used where the top 30 most intense ions were chosen for MS/MS fragmentation. Raw data were then processed using MaxQuant, which implements the Andromeda search engine, for peptide and protein identification and SILAC quantitation.

For the proteomics dataset 1155 out of 2024 probes had a significant *p*-value<0.05 (calculated in MaxQuant). To identify differentially expressed proteins between time 0 and 4, 8 and 12h we collected per time point the proteins whose absolute ratios were >0.5. For each time point, the proteins were then split whether the ratio was positive or negative. 6 lists were annotated for GO terminology using GeneSpring. The extensive list of GO-terms was submitted to Revigo (Supek et al., 2011) to reduce complexity and the resulting GO-categories depicted in Extended Data Fig. 1f. The mass spectrometry proteomics data have been deposited with the ProteomeXchange Consortium via the PRIDE partner repository with the dataset identifier PXD003281.

### siRNA nucleofection

siRNA nucleofection was performed with the Amaxa 16-well shuttle system (Lonza). Pre-confluent keratinocytes were disaggregated and resuspended in cell line buffer SF. For each 20μl transfection (program FF-113), 2 × 10^5^ cells were mixed with 1–2 μΜ siRNA duplexes as described previously (Mulder et al., 2012). Transfected cells were incubated at ambient temperature for 5–10 min and subsequently replated in pre-warmed Keratinocyte-SFM medium until required for the downstream assay. SMART pool ON-TARGET plus siRNAs (Ambion/GE Healthcare) were used for gene knockdowns. Each SMART pool was a mix of 4 sets of RNAi oligos. The sequences of the siRNA oligos are provided in the Extended Data Table 11.

### Doxycycline and Cumate inducible overexpression

For the Doxycycline inducible system we used the pCW57-GFP-2A-MCS lentiviral vector (gift from Adam Karpf; Addgene plasmid # 71783). Primary keratinocytes were transduced with lentiviral particles containing protein expression vector encoding genes for wild type DUSP6^C293S^, mutant DUSP6 and DUSP10^C293S^. 2 days post-transduction, cells were subjected to Puromycin selection for 3 days and then 1μg/ml Doxycycline was added to the growth medium.

For Cumate induction we used the lentiviral QM812B-1 expression vector (System Biosciences). Cells were transduced and selected as described for the Doxycycline system and protein expression was induced by addition of 30 μg/ml Cumate solution to the growth medium.

### Clonogenicity assays

100, 500 or 1000 keratinocytes were plated on a 3T3 feeder layer per 10cm^2^ dish or per well of a 6 well dish. After 12 days feeders were removed and keratinocyte colonies were fixed in 10% formalin (Sigma) for 10 min then stained with 1% Rhodanile Blue (1:1 mixture of Rhodamine B and Nile Blue A (Acros Organics). Colonies were scored by ImageJ and clonogenicity was calculated as the percentage of plated cells that formed colonies.

### Skin reconstitution assays

Pre-confluent keratinocyte cultures (km passage 3) were disaggregated and transfected either with SMART pool siRNAs or non-targeting control siRNAs. 24 hours post-transfection, keratinocytes were collected and reseeded on irradiated de-epidermised human dermis (Sen et al., 2010) in 6-well Transwell plates with feeders and cultured at the air–liquid interface for three weeks. Organotypic cultures were fixed in 10% neutral buffered formalin (overnight), paraffin embedded and sectioned for histological staining. 6μm thick sections were labelled with haematoxylin and eosin or appropriate antibodies. Images were acquired with a Hamamatsu slide scanner and analysed using NanoZoomer software (Hamamatsu).

### Epidermal wholemounts

The procedure was modified from previous reports (Jensen et al., 1999). Skin samples from either breast or abdomen were obtained as surgical waste with appropriate ethical approval and treated with Dispase (Corning) overnight on ice at 4^°^C. The epidermis was peeled off as an intact sheet and immediately fixed in 4% paraformaldehyde for 1 hour. Fixed epidermal sheets were washed and stored in PBS containing 0.2% sodium azide at 4°C. Sheets were stained with specific antibodies in a 24-well tissue culture plate. Image acquisition was performed using a Nikon A1 confocal microscope. 3D maximal projection (1,024 × 1,024 dpi), volume rendering and deconvolution on stacked images were generated using NIS Elements version 4.00.04 (Nikon Instruments Inc.).

### Western blotting

Cells were lysed on ice in 1x RIPA buffer (Bio-Rad) supplemented with protease and phosphatase inhibitor cocktails (Roche). Total protein was quantified using a BCA kit (Pierce). Soluble proteins were resolved by SDS-PAGE on 4-15% Mini-PROTEAN TGX gels (Bio-Rad) and transferred onto Trans-Blot 0.2um PVDF membranes (Bio-Rad) using the Trans-Blot Turbo transfer system (Bio-Rad). Primary antibody probed blots were visualized with appropriate horseradish peroxidase-coupled secondary antibodies using enhanced chemiluminescence (ECL; Amersham). The ChemiDoc Touch Imaging System (Bio-Rad) was used to image the blots. Quantification of detected bands was performed using Image Lab software (Bio-Rad).

### Antibodies

Antibodies against the following proteins were used: P-ERK (Cell Signaling # 9101; western blot – 1:1000 dilution), ERK (Cell Signaling # 9102; western blot – 1:1000), P-p38 (Cell Signaling # 9211; western blot – 1:1000), p38 (Cell Signaling # 9212; western blot – 1:1000), α-Tubulin (Sigma # T6199; western blot – 1:5000), MKP-3/DUSP6 (Abcam # ab76310; western blot – 1:1000 and R&D Systems *#* MAB3576-SP; immunostaining – 1:200), PPTC7 (Abcam # ab122548; western-blot 1:250 and Sigma # HPA039335; immunostaining – 1:200), PTPN1/PTP1B (Sigma # HPA012542; western-blot – 1:500; R&D Systems # AF1366-SP; immunostaining – 1:200), PTPN13 (R&D Systems # AF3577; western-blot - 1:300 and immunostaining – 1:200), PPP3CA/Calcineurin A (Sigma # HPA012778; western-blot 1/1000 and R&D Systems # MAB2839-SP; immunostaining – 1:200), DUSP10 (Abcam # 140123; western-blot - 1:1000 and immunostaining – 1:200), TP63 (SCBT # sc367333; immunostaining – 1:100), Involucrin (SY5, in-house; immunostaining – 1:500) and β1-Integrin (clone P5D2, in-house; immunostaining – 1: 200). Species-specific secondary antibodies conjugated to Alexa 488 or Alexa 594 were purchased from Molecular Probes, and HRP-conjugated second antibodies were purchased from Amersham and Jackson ImmunoResearch.

### RNA extraction and RT–qPCR

Total RNA was isolated using the RNeasy kit (Qiagen). Complementary DNA was generated using the QuantiTect Reverse Transcription kit (Qiagen). qPCR analysis of cDNA was performed using qPCR primers (published or designed with Primer3) and Fast SYBR green Master Mix (Life Technologies). RT-qPCR reactions were run on CFX384 Real-Time System (Bio-Rad). Heatmaps of RT-qPCR data were generated by Multiple Expression Viewer (MeV_4_8) or GraphPad Prism 7. Sequences of qPCR primers are provided in Extended Data Table 12.

### Protein phosphatase networks

The mechanistic networks depicted in Fig. 4a were built with Cytoscape (cytoscape.org, V3.2.1). The data from Extended Data Table 1 were used to decide whether or not there was a statistically significant interaction between any two phosphatases (adjusted p-value <0.05) and the fold-change in the Table gives the directionality of the edge. The node size shown in Fig. 4a is proportional to the phosphatase expression at that time point.

### Statistics

Hierarchical clustering of the genomics and proteomics data was generated either by MeV_4_8 or MatLab. Statistical analysis of the quantifications presented in the Figure legends was performed using GraphPad Prism 7.

## Acknowledgements

This research was funded by the Medical Research Council and the Wellcome Trust. The authors also gratefully acknowledge the use of Core Facilities provided by the financial support from the Department of Health via the National Institute for Health Research (NIHR) comprehensive Biomedical Research Centre award to Guy’s & St Thomas’ NHS Foundation Trust in partnership with King’s College London and King’s College Hospital NHS Foundation Trust. Benedicte Oules is the recipient of fellowships from Societe Francaise de Dermatologie, Collège des Enseignants en Dermatologie de France and Fondation d’Entreprise Groupe Pasteur Mutualité. We thank Giacomo Donati, Arsham Ghahramani and Klaas Mulder for helpful advice and discussions and are grateful to Fredrik Pontén for providing antibodies.

## Author contributions

FMW and AIL conceived the project and oversaw the experiments. AM, AOP, BO, TL, K LA, PV, JN and GW performed and analysed experiments. AOP and S-J D performed computational analysis. AOP performed bioinformatic analysis. All authors contributed to preparing the paper.

## Author information

The authors have no conflicts of interest to declare.

---

**Extended Data Figure 1.**
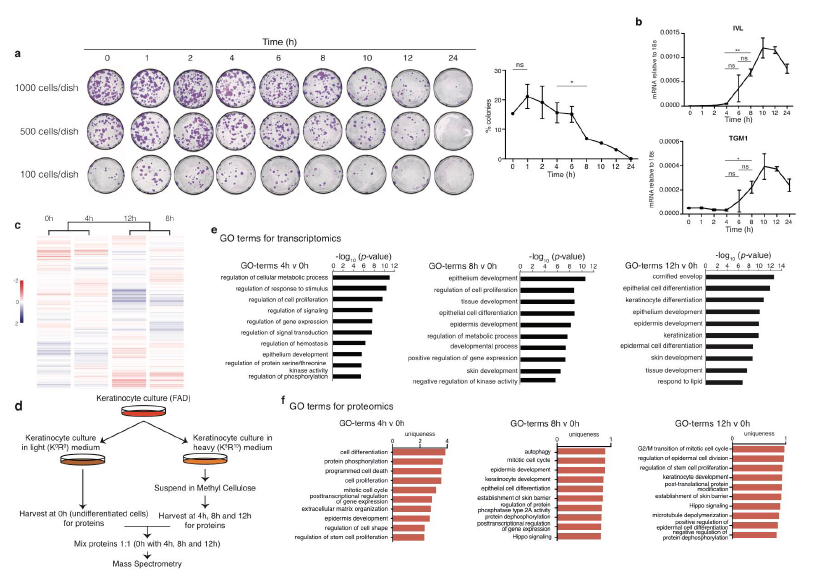
Clonal growth, genomic and proteomic analysis of suspension-induced terminal differentiation. **a.** Keratinocytes harvested after 0, 1, 2, 4, 6, 8, 10, 12 or 24h in suspension in methylcellulose were seeded at 100, 500 or 1000 cells in 10 cm^2^ culture dish on mitotically inactivated J2 3T3 feeders and cultured for 12 days. Following staining, colonies were counted using ImageJ (*n*=*2* independent experiments with 3 replicate dishes per experiment; *p*-value calculated by Tukey’s multiple comparison test). **b.** RT-qPCR quantification of IVL and TGM1 mRNA levels at different times in suspension (*n*=3 independent cultures; two tailed *t*-test). **c.** Hierarchical clustering of significantly expressed transcripts at 0h, 4h, 8h and 12h in suspension (each time point represents the mean value of n=3 independent experiments). **d.** Schematic of SILAC-Mass Spectrometry labelling protocol. **e.** GO analysis of differentially expressed genes upregulated at 4, 8 and 12h relative to 0h. The bar plots represent −log_10_ of *p*-values of the identified GO terms. **f.** GO analysis of proteins ranked in the order of their expression level (fold increase relative to 0h) at 4, 8 and 12h. GO terms were fetched for individual proteins rather than for clusters of proteins. Bar plots represent −log_10_ of the *p*-values of the identified GO terms. Error bars represent s.d. * *p* < 0.05; *\*\*p* < 0.01; ns = non-significant.

**Extended Data Figure 2.**
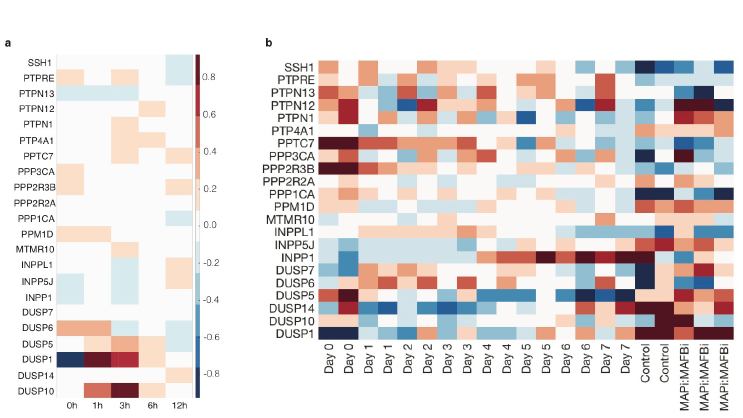
Expression in published datasets of the protein phosphatases identified by suspension-induced differentiation. **a.** Heatmap showing mRNA expression of candidate phosphatases during calcium-induced stratification of primary human keratinocytes (GSE38628). **b.** Heatmap showing mRNA expression of candidate phosphatases during ex vivo human epidermal reconstitution (GSE52651).

**Extended Data Figure 3.**
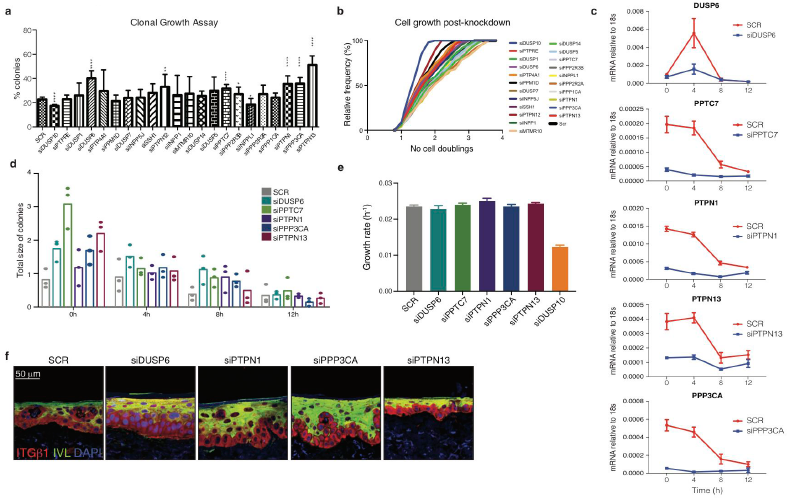
Effects of phosphatase knockdown on keratinocyte growth and differentiation. **a.** Effect of phosphatase knockdown on colony formation. Values plotted are average % colony formation in three independent screens with three replicate dishes per screen. Error bars represent s.d. *p*-values were generated using two tailed *t*-test (\**p* < 0.05; \*\**p* < 0.01; \*\*\*\**p* < 0.0001; ns = non-significant). **b.** Following siRNA transfection keratinocytes were seeded in collagen-coated 96 well format dishes and cultured in KSFM medium for three days. 24h post transfection the culture dishes were moved into an Incucyte microscope and scanned every hour for the next 48 hours to evaluate cell growth and proliferation. Each data point is the average of 3 replicate screens with 8 independent well scans per condition per screen. Values at each time point were normalized to the respective 0h time point or the first scan point. A cumulative plot is shown with the relative frequency (%) of cell doublings. **c.** RT qPCR quantification of phosphatase mRNA levels relative to 18s RNA as a function of time in suspension, showing efficiency of the different knockdowns in the starting populations. Error bars represent s.d. **d.** Effect of phosphatase knockdowns on the total area of colonies per dish after cells were replated after different times in suspension Data are expressed to SCR control. Mean % areas and individual values are plotted (*n*=*3* independent transfections). **e.** Growth rate of human keratinocytes after phosphatase knockdown, calculated by fitting the data in (**b**) to an exponential growth curve and averaging the rates. **f.** Sections of reconstituted epidermis labelled with antibodies to ITGβ1 and IVL with DAPI as nuclear counterstain. Scale bar: 50 μm.

**Extended Data Figure 4.**
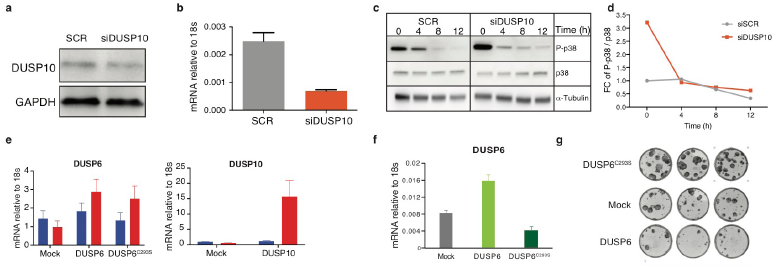
Effects of DUSP10 knockdown and DUSP6 and DUSP10 overexpression. **a, b.** Western blot (**a**) and RT qPCR measurements (**b**) of DUSP10 knockdown by SMART pool siRNA. **c, d.** Western blot of phospho-p38 and total p38 in suspended cells following DUSP10 knockdown. siSCR, α-tubulin loading control and quantitation are shown. **e.** RT qPCR quantification of doxycycline-induced over-expression of DUSP6, mutant DUSP6^C293S^ and DUSP10 relative to 18s mRNA. Cells were treated with 1μg/ml doxycycline for 24h. **f.** RT qPCR quantification of cumate-induced over-expression of DUSP6 and mutant DUSP6^C293S^ relative to 18s mRNA. **g.** Representative dishes showing effects of overexpressing wild type and mutant DUSP6 on clonal growth (representative of *n*=3 dishes).

**Extended Data Figure 5.**
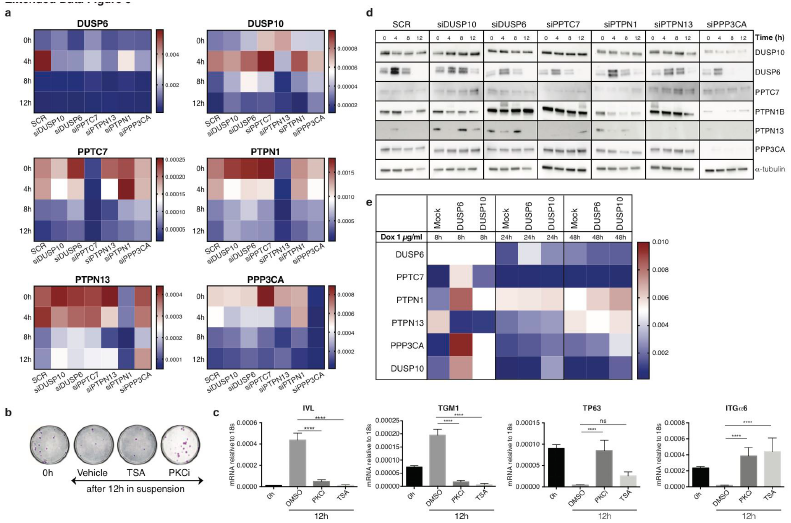
Effects on phosphatase expression of knockdowns and treatment with TSA or PKCi. **a.** Effect of TSA and PKCi on clonogenicity of keratinocytes following suspension in methylcellulose for 12h. **b.** mRNA levels of IVL, TGM, ITGα6, and TP63 in cells held in suspension for 12h. Cells were treated with TSA, PKCi or DMSO (vehicle control) (*n* = 3 independent treated cultures with two technical replicates each). *p-*values for the comparisons were generated by Tukey’s multiple comparison test. **c.** Heatmaps showing the effect of knocking down individual phosphatases on mRNA levels of other phosphatases with time in suspension (0h, 4h, 8h, 12h). RT-qPCR is relative to 18s mRNA (*n*=3 independent transfections; see Extended Data Table 8 for *p*-values generated by twoway ANOVA with Dunnett’s multiple comparisons test). **d.** Western blots showing phosphatase levels in primary keratinocytes upon knockdown of scrambled control (SCR), DUSP6, PPTC7, PTPN1, PTPN13 or PPP3CA and suspension for 0, 4, 8 or 12h. α-tubulin: loading control. **e.** RT qPCR quantification of phosphatase mRNA levels (relative to 18s mRNA) following doxycycline-induced over-expression of DUSP6, mutant DUSP6^C293S^ and DUSP10. Cells were treated with 1μg/ml doxycycline for 8h, 24h or 48h (*n*=3 independent cultures; see Extended Data Table 9 for *p*-values generated by two-way ANOVA with Dunnett’s multiple comparisons test).

## Extended Data Tables

**Extended Data Table 1** Log_2_ fold change of normalized gene expression for all pairwise comparisons of mRNA levels during suspension-induced terminal differentiation. For each condition the mean of n=3 independent replicates was used and the pairwise fold change comparison is between the means of both samples.

**Extended Data Table 2:** Proteomics data for all pairwise comparisons of protein levels at 4h, 8h and 12h in suspension relative to the 0h control. For each condition the mean of n=3 independent replicates was used and the pairwise fold change comparison is between the means of both samples.

**Extended Data Table 3:** Phosphoproteomics data for pairwise comparisons at 4h and 8h in suspension relative to the 0h control.

**Extended Data Table 4:** Log2 ratio of phosphopeptides over total proteins at 4h.

**Extended Data Table 5:** p-values generated for RT qPCR of TP63 and TGM1 for each conditional time course relative to control time course (SCR) by 2-way ANOVA with Dunnett’s multiple comparisons test (related to Fig. 2f, g).

**Extended Data Table 6:** Effect of phosphatase knockdown on AP1 transcription factor expression. *p*-values generated for each conditional time course relative to control time course (SCR) by 2-way ANOVA multiple comparisons (for AP1 superfamily factors). p-values generated for RT qPCR of AP1 factors for each conditional time course relative to control time course (SCR) by 2-way ANOVA with Dunnett’s multiple comparisons test.

**Extended Data Table 7:** Effect of DUSP6 and DUSP10 knockdown on AP1 transcription factor expression. *p*-values generated for RT qPCR of AP1 factors relative to control cells (SCR) by 2-way ANOVA.

**Extended Data Table 8:** p-values generated for RT qPCR of phosphatases for each conditional time course relative to control time course (SCR) by 2-way ANOVA with Dunnett’s multiple comparisons test.

**Extended Data Table 9:** One-way non-parametric ANOVA (Friedman test) with Dunn’s multiple comparisons test for the effect of overexpressing DUSP6 and DUSP10 on mRNA levels of the pro-commitment phosphatases, determined by RT-qPCR.

**Extended Data Table 10**: Boolean expression patterns and phosphatases interactions used to generate Fig. 4c, d.

**Extended Data Table 11**: siRNA library for phosphatase knockdown.

Extended Data Table 12: **List of qPCR primers.**

